# Broad-scale puma connectivity could restore genomic diversity to fine-scale coastal populations

**DOI:** 10.1101/2021.10.08.463677

**Authors:** Kyle D. Gustafson, Roderick B. Gagne, Michael R. Buchalski, T. Winston Vickers, Seth P.D. Riley, Jeff Sikich, Jaime L. Rudd, Justin A. Dellinger, Melanie E.F. LaCava, Holly B. Ernest

## Abstract

Urbanization is decreasing wildlife habitat and connectivity worldwide, including for apex predators, such as the puma (*Puma concolor*). Puma populations along California’s central and southern coastal habitats have experienced rapid fragmentation from development, leading to calls for demographic and genetic management. To address urgent conservation genomic concerns, we used double-digest restriction-site associated DNA (ddRAD) sequencing to analyze 16,285 genome-wide single-nucleotide polymorphisms (SNPs) from 401 broadly sampled pumas. Our analyses indicated support for 4–10 geographically nested, broad- to fine-scale genetic clusters. At the broadest scale, the 4 genetic clusters had high genetic diversity and exhibited low linkage disequilibrium, indicating pumas have retained statewide genomic diversity. However, multiple lines of evidence indicated substructure, including 10 fine-scale genetic clusters, some of which exhibited allelic fixation and linkage disequilibrium. Fragmented populations along the Southern Coast and Central Coast had particularly low genetic diversity and strong linkage disequilibrium, indicating genetic drift and close inbreeding. Our results demonstrate that genetically at-risk populations are typically nested within a broader-scale group of interconnected populations that collectively retains high genetic diversity and heterogeneous fixations. Thus, extant variation at the broader scale has potential to restore diversity to local populations if management actions can enhance vital gene flow and recombine locally sequestered genetic diversity. These state- and genome-wide results are critically important for science-based conservation and management practices. Our broad- and fine-scale population genomic analysis highlights the information that can be gained from population genomic studies aiming to provide guidance for fragmented population conservation management.

## INTRODUCTION

Human development is reducing habitat on a global scale, undermining efforts to conserve ecosystem structure and function (Newbold et al., 2016). Reports of fragmented wildlife populations and the increasing need for human housing and associated agriculture and energy have emphasized the necessity for development to avoid impacting the long-term sustainability of wildlife populations (Jordan et al., 2007; Kiesecker et al., 2011; Saha & Paterson, 2008). One of the most developed states in the United States is California, which contains the largest census size with over 39 million people (US Census, 2019). Although the development of California has led to historical extirpations of other apex predators, such as the grizzly bear (*Ursus arctos*; Herrero, 1970) and gray wolf (*Canis lupus*; Schmidt, 1991), the puma (*Puma concolor;* also known as mountain lion and cougar) has maintained a widespread distribution throughout the state (Dellinger et al., 2020a).

Currently, approximately half of all apparent puma habitat in California is conserved and the remainder could be subject to further development (Dellinger et al. 2020a). Much of the inland areas of California have continous streches of protected habitat (Dellinger et al. 2020a) supporting puma populations with high genetic diversity and large effective populations sizes (Gustafson et al., 2019). However, movement corridors among coastal mountain ranges are increasingly being degraded by human development (Burdett et al., 2010; Suraci, Nickel, & Wilmers, 2020; Zeller et al., 2017). Despite the natural long-range dispersal abilities of pumas (Gonzalez-Borrajo, López-Bao, & Palomares, 2017), interstate highways limit dispersal via avoidance and direct mortality in some urban areas (Riley et al., 2014; Vickers et al., 2015). Although human-caused mortality from vehicle collisions and lethal removal after wildlife– livestock conflicts are concerns (Guerisoli, Luengos Vidal, Caruso, Giordano, & Lucherini, 2020; Torres, Mansfield, Foley, Lupo, & Brinkhaus, 1996), a larger concern for long-term population viability is the genetic isolation of pumas within small or shrinking patches of habitat, which has led to high levels of intraspecific competition and mortality (Benson, Sikich, & Riley, 2020) and low genetic diversity in some areas (Ernest et al., 2014; Gustafson et al. 2019; Riley et al., 2014).

Previous studies have reported that two isolated puma populations in southern California, including the Santa Ana Mountains and the Santa Monica Mountains (**Fig. 1**), had the lowest genetic diversity estimates measured (Ernest et al., 2014; Riley et al., 2014), apart from the endangered Florida panther (*P. c. coryi*). In the Santa Ana Mountains, phenotypic evidence of inbreeding depression in the form of kinked tails has been observed, similar to Florida panthers (Ernest et al., 2014; Roelke, Martenson, & O’Brien, 1993). For both populations, freeway traffic is isolating pumas (Ernest et al., 2014; Riley et al., 2014; Vickers et al., 2015) and contemporary gene flow has been severely limited. Detailed pedigree analyses following the immigration of one male into each region showed evidence of natural genetic rescue (Ernest et al., 2014; Gustafson, Vickers, Boyce, & Ernest, 2017; Riley et al., 2014). Although migrant effects were positive, projection models predict the extirpation of these populations in 50 years without enhanced demographic dispersal and gene flow (Benson et al., 2016; 2019).

**Figure 1.**
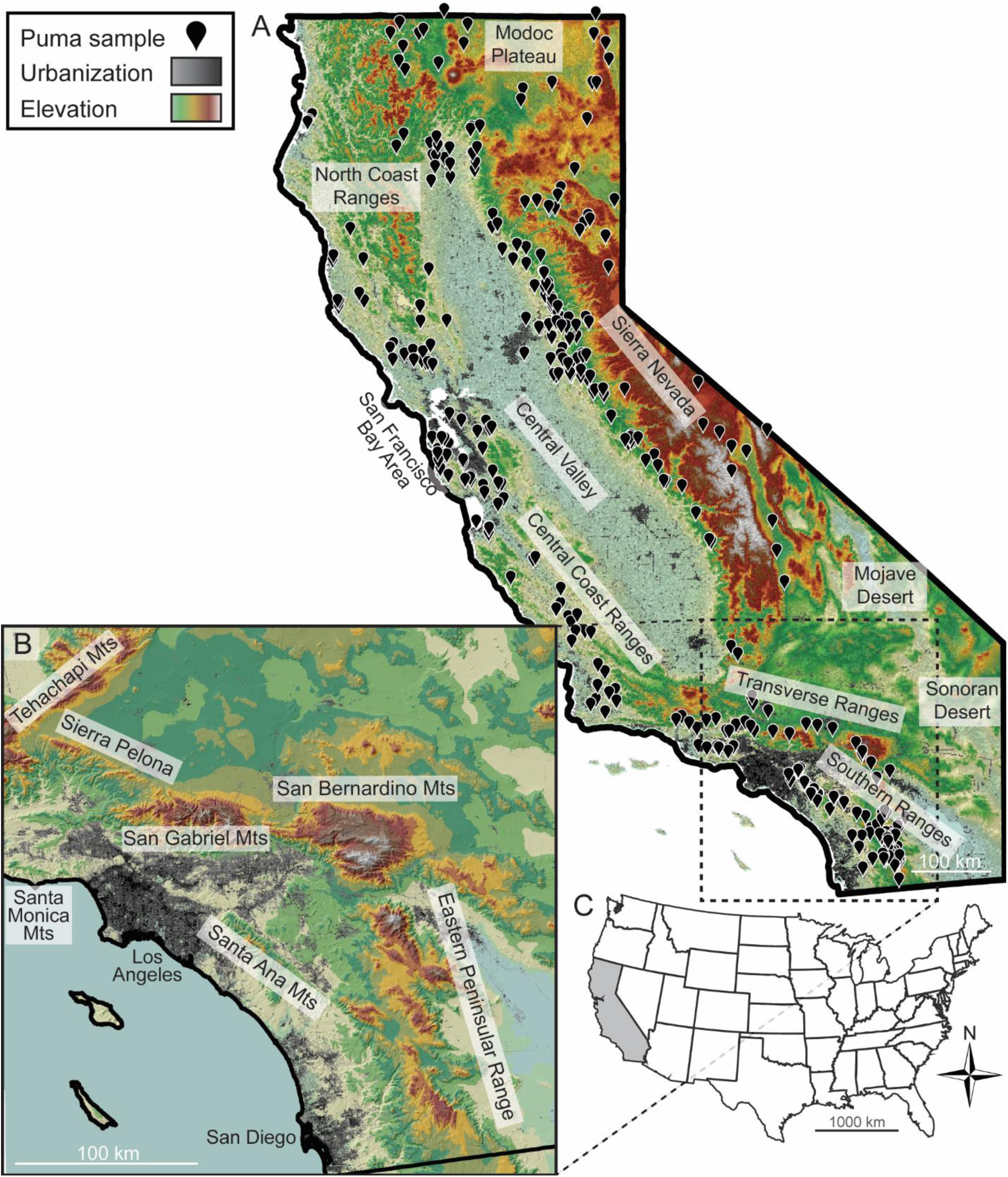
Location of 401 sampled pumas used in analyses, including (**A**) sample distribution across California, (**B**) geography of the mountain ranges surrounding the Los Angeles and San Diego regions, and (**C**) inset map showing the location of California in the United States of America.

Recently published genome-resequencing data that included 4 pumas from California; 2 pumas from the Santa Monica Mountains and 2 pumas from the Central Coast North region in the Santa Cruz Mountains; indicated pumas from those coastal ranges had ∼20–40% of their genomes represented as long runs of homozygosity, resulting from recent inbreeding (Saremi et al., 2019). However, these runs of homozygosity were not shared among individuals, and different populations exhibited different homozygous haplotypes, suggesting genetic restoration (Hedrick, 2005; Tallmon, Luikart, & Waples, 2004) is possible because genetic variation still exists.

The complex distribution of pumas throughout California along a continuum of high genetic diversity populations occupying abundant habitat, to strongly isolated populations displaying evidence of inbreeding depression, requires a thorough characterization of statewide genomic diversity to achieve proper conservation. In this study, our objective was to characterize patterns of genomic diversity at varying geographic scales. Such an approach has the potential to aid conservation strategies because it can identify at-risk, low-diversity local populations that would benefit from restored gene flow within a broader geographic region. We identified single-nucleotide polymorphisms (SNPs) from hundreds of individuals using a double-digest, restriction-site associated DNA sequencing method (ddRAD; Peterson, Weber, Kay, Fisher, & Hoekstra, 2012). Specifically, we assessed population structure, genetic diversity, evidence for selection, and linkage disequilibrium using 16,285 SNPs among 401 pumas. Our study provides a contemporary analysis of puma population genomics and new insights that can guide management actions to restore connectivity among puma populations throughout the state of California.

## METHODS

### Sample collection and DNA extraction

We obtained 354 muscle tissue samples collected by the California Department of Fish and Wildlife between 2011–2017 from pumas either hit-by-car (∼6%), found dead (∼2%), poached (<1%), or through depredation permits (>90%) which had never been used in any previous genetic survey. Samples were well-distributed throughout the state, except for smaller populations in smaller mountain ranges. To bolster our sample size in the Los Angeles region of southern California, we added 144 pre-existing DNA extracts from pumas collected between 2002–2015 (Riley et al. 2014; Vickers et al. 2015). After genomic and bioinformatic filtering (described below), we retained 401 out of 498 samples in the final dataset, which spanned the majority of puma habitat in California, excluding desert regions (**Fig. 1**). For samples that lacked a precise GPS location, we used the nearest address or town where they were collected as their GPS point. Samples were stored at -80°C until DNA was extracted DNA using Omega Bio-tek Mag-Bind Blood & Tissue DNA HDQ Kits (Omega Bio-tek, #M6399-01) with a manufacturer-designed protocol for the Kingfisher Duo Prime (ThermoFisher Scientific, #5400110) automated DNA purification system. We measured the concentration of DNA from each sample using a Qubit 3.0 fluorometer (Invitrogen, #Q33216) with Qubit dsDNA high-sensitivity kits (Invitrogen, #Q32854).

### Double digest restriction site associated DNA library preparation and sequencing

We reduced the genome size of our samples and identified single-nucleotide polymorphisms (SNPs) using modifications to the double digest restriction-site associated DNA sequencing (ddRAD) protocols developed by Peterson et al. (2012). We used a library construction scheme which pooled 48 samples per library based on barcode availability, cost effective multiplexing, and sufficient coverage per individual. For each pooled library, we first normalized DNA concentrations to the sample with the lowest concentration within a library, with the goal to be above 200ng DNA starting material in 25μL elution buffer (>8 ng/μL). The library with the lowest normalized starting concentration for each sample had 17.8 ng/μL DNA, whereas the library with the highest starting material had 51.6 ng/μL DNA. We used digestion enzymes and protocols established with previous puma work (Trumbo et al., 2019). After DNA was normalized, we double-digested the DNA from each individual using NlaIII (New England BioLabs, #R0125S) and EcoRI (New England BioLabs, #R3101S) restriction enzymes (37°C for 3 hours, then held at 4°C) at manufacturer-recommended enzyme concentrations and used AMPure XP beads (Beckman Coulter, #A63881) at a 1.5X ratio to retain only DNA from the digestion. We omitted the DynaBeads cleanup and again used the Qubit to measure DNA concentrations and to guide another round of normalization. After normalization, the library with the lowest per-sample concentration had 2.1 ng/μL (in 29 μL) and the library with the highest per-sample concentration had 8.1 ng/μL.

We then ligated 48 uniquely barcoded P1 adaptors (e.g., P1.1 through P1.48) and two common P2 adapter pairs (i.e., P2.1 and P2.2) to each sample’s double-digested fragments using the protocols of Peterson et al. (2012) to identify individual puma samples. Following ligation with individual barcodes, we pooled all 48 samples into a single tube and used AMPure XP beads to clean the library. We used TE buffer (rather than molecular-grade water) as the final step in this cleanup, which is recommended by the manufacturer for running size selection in the Pippin Prep (Sage Science, Beverly, Massachusetts). We selected fragments ranging from 375–475 bp (including 75bp of adapters) using 2% dye-free gels run on a Pippin Prep. To minimize random PCR duplicate errors, we split the library and ran 5 high-fidelity Phusion (New England BioLabs, #M0530) PCRs for 12 cycles on a SimpliAmp thermal cycler (ThermoFisher Scientific, #A24811). We then recombined the 5 PCR products and used an AMPure XP bead clean up on the amplified library. Sample concentrations after size selection averaged 2.0 ng/μL DNA (range 0.82–3.7) and, after the PCR, averaged 8.2 ng/μL (range 3.6–15.0). We shipped the unfrozen DNA with freezer packs to the University of Oregon’s Genomics and Cell Characterization Core Facility (https://gc3f.uoregon.edu/) for 150bp single-end sequencing on an Illumina HiSeq 4000 (Illumina, San Diego, CA).

### Bioinformatic SNP filtering

We ran standard quality control analyses using program *FastQC* v0.11.5 (Andrews, 2010). We used the *process_radtags* program in the *Stacks* v2.55 (Catchen, Hohenlohe, Bassham, Amores, & Cresko, 2013) package to de-multiplex the reads based on unique barcodes, to assign each sequence to an individual puma sample, to remove sequences with a Phred quality score below 20 (99% accuracy), and to remove Illumina adapter sequences from the data. We then aligned reads for each individual to PumCon1.0—the *Puma concolor* draft reference genome —using program *bwa* (Li & Durbin, 2009). We identified and filtered SNPs with *Samtools* (Li et al., 2009). We discarded loci with a mapping quality score below 20, minimum base quality less than 20, with more than two alleles at a site, and with a maximum depth greater than 100. We skipped indels and only used a random SNP per read to reduce linkage disequilibrium.

Using *vcftools*, we tested the effects of multiple filtering parameters on our dataset, specifically looking at which parameters produced unreliable and inconsistent heterozygosity estimates, inbreeding coefficients, and relatedness values. We retained loci with a minor allele frequency ≥ 0.05 as lower frequency SNPs could be sequencing error. The relationship between minimum depth of reads per individual and heterozygosity was asymptotic and plateaued at about 3–4 reads. To be conservative, we selected a minimum depth of 4 reads per individual to reliably acquire genotypes based on both alleles. We also retained SNPs that had genotypes for at least 50% of the individuals. We iteratively removed samples with more than 50% missing data to maximize the number of SNPs retained in the dataset. Being more conservative with the percent of missing data only decreased the number of SNPs in the final dataset and did not affect heterozygosity estimates, inbreeding coefficients, and relatedness values. We scanned for duplicate samples using relatedness values in *vcftools*, but, as expected, found none because all DNA samples were removed from dead pumas. We also removed two potentially contaminated samples based on negative F statistics in *vcftools*.

In each library of 48 samples, we strategically included puma samples from across a large geographic area so libraries would have no correlation with spatial location. For example, there was no significant difference between mean sample latitudes (*F*_7;309_ = 1.108, *p* = 0.358) or longitudes (*F*_7;309_ = 1.533, *p* = 0.155) among libraries. However, because the southern California libraries constructed from pre-existing extracts were from a small geographic region, there ended up being some latitudinal (*F*_10;395_ = 33.76, *p* < 0.001) and longitudinal (*F*_10;395_ = 33.89, *p* < 0.001) mean differences between those libraries and the libraries constructed from the new samples.

To test for library effects (i.e., differences among sequencing lanes), we used *BayeScan* to identify outlier SNPs while treating sequencing lanes as “populations” and using a false discovery rate of 0.05 (Foll & Gaggiotti, 2008). We also assessed bias with various genetic structure analyses (see below). Genotypes resulting from the pre-existing DNA extracts consistently clustered with those genotypes resulting from the new samples collected from southern California. With no apparent library effects, we retained 16,285 bi-allelic variants (mean ± SD = 12,245 ± 2749) with a mean depth at each locus of 11.7 ± 5.1 and a mean depth per locus per individual of 11.7 ± 7.1.

### Population structure and outlier loci

We used multiple approaches to identify genetic clusters of individuals, including a linear principal components analysis (PCA) and a spatially-explicit population structure analysis in program R. We ran the PCA using *adegenet* 2.1.1 (Jombart, 2008) and the structure analysis in *tess3r* 1.1.0 (Caye, Deist, Martins, Michel, & François 2016). We used *adegenet::colorplot* to present linear structure identified by the first 3 principal component axes. In *tess3r*, we ran 20 replicates for each *K* (1–20) at 100,000 iterations each. We kept the most highly supported model (i.e., “best” based on cross-entropy scores) within each of the 20 replicates. To test for evidence of loci under selection, we identified outlier loci among populations (Narum & Hess, 2011) using *BayeScan* and *tess3r* with the Benjamini–Hochberg statistical correction and the recommended α-value of 0.0001.

### Genetic diversity, effective population size, genetic differentiation, and linkage decay

For each genetic cluster identified in *tess3r*, we calculated observed heterozygosity (*H*_*O*_), gene diversity (*H*_*S*_), and allelic richness (*A*_*r*_) using *hierfstat::basic*.*stats* (Goudet, 2005). To test for Wahlund effects within broad-scale clusters, we used t-tests to test for differences between *H*_*O*_ and *H*_*S*_. We calculated private alleles (*A*_*p*_) using *poppr::private_*alleles (Kamvar, Tabima, & Grünwald, 2014). We used *NeEstimator* 2.1 (Do et al., 2014) to estimate effective population size (*N*_e_), using the linkage disequilibrium model with allele frequencies >0.05 with a correction factor of 19 haploid chromosomes (Hsu, Rearden, & Luquette, 1963) as recommended by (Waples, Larson, & Waples, 2016). We used *hierfstat::pairwise*.*neifst* and *hierfstat::pairwise*.*WCfst* to estimate pairwise genetic differentiation based on *F*_*ST*_ according to Nei (1987) or Weir and Cockerham (1984).

We used *Plink* 2.0 (Purcell et al., 2007) to estimate linkage disequilibrium among loci (--ld-window-r2 0 --ld-window 999999 --ld-window-kb 8000). To determine the level of non-random segregation of alleles across the genome, we assessed linkage decay in each genetic cluster by plotting the correlation of loci (*R*^*2*^) based on genomic distance between SNPs. We correlated loci using binned intervals of 100,000bp from 0 to the maximum scaffold size of PumCon1.0. Meiosis should break up linkage, resulting in low *R*^*2*^ values. However, populations experiencing strong selection, low mutation, inbreeding, low migration, or strong genetic drift will have higher *R*^*2*^ values. In short, SNPs that are close together on chromosomes are expected to be correlated (i.e., inherited as chromosomal/haplotype segments), but SNPs far away are expected to assort randomly during recombination. However, if sequences are too similar, which they may be in small and inbred populations, we are not be able to detect events of crossing over despite their occurrence, resulting in higher estimates of linkage disequilibrium, which is still an important indicator of genetic diversity and *N*_e_.

## RESULTS

### Population structure and outlier loci

We recovered 16,285 SNPs. The first three axes of the PCA (i.e., linear combinations of SNPs) accounted for 14.6% of the variance and indicated there were 4 broad-scale genetic clusters distributed across California (**Fig. 2**). When each puma was plotted on a map of California (**Fig. 2A**), the 4 clusters were geographically concordant with the Sierra Nevada (SN), North Coast (NC), Central Coast (CC), and Southern Coast (SC). The first eigenvector separated the negative-valued CC and SC groups from the positive-valued SN and NC (**Figs 2B & 2C**). The second eigenvector separated negative-valued CC from positive-valued SC (**Fig. 2B**). Finally, the third eigenvector separated negative-valued NC from all other groups (**Fig. 2C**).

**Figure 2.**
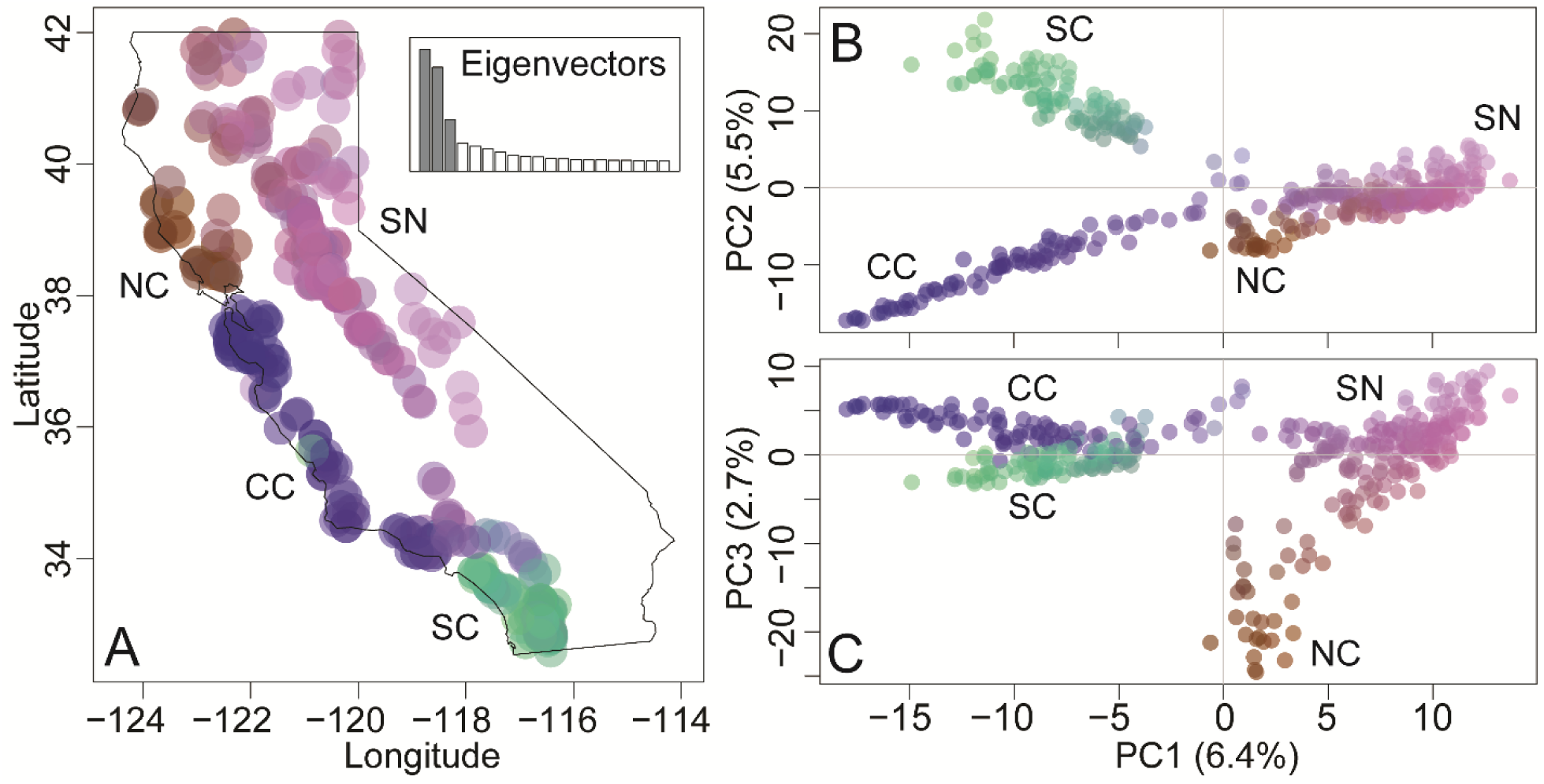
Principal component analysis (PCA) of 401 pumas at 16,285 SNPs reveals four genetic clusters. **(A)** The colorplot (R package *adegenet*) of the PCA represents colors corresponding to a combination of the first 3 eigenvectors. The inset plot shows the proportion of the variance explained by shaded PC eigenvectors 1–3 compared to other eigenvectors. The color values are plotted at sample locations to demonstrate geographic structure. Colorplots of **(B)** PC1 and PC2 and **(C)** PC1 and PC3 resolved the 4 broad-scale genetic clusters.

A spatially-explicit population structure analysis indicated that there was broad- to fine-scale nested genetic structure with support for 4–10 genetic clusters (**Fig. 3**). Root mean square error (inset plot in *K* = 2 panel of **Fig. 3**) and cross-entropy scores (inset plot in *K* = 3 panel of **Fig. 3**) provide statistical evidence for nested genetic structure; values begin to plateau at *K* = 4 but there is a steady increase in statistical support at higher *K* values. However, single pumas formed individual clusters at *K* > 10 at which point *K* lost biological meaning. When *K* was set to 4, the genetic clusters corresponded to the broad-scale genetic groups identified by the PCA (**Figs. 2 & 3**). Briefly, at *K* = 5, pumas in the Central Coast North (CC-N) emerged; at *K* = 6, the Eastern Sierra Nevada (ESN) cluster separated from the Western Sierra Nevada (WSN); at *K* = 7, the Santa Ana (SA) cluster separated from the Eastern Peninsular Range (EP); at *K* = 8, the San Gabriel–San Bernardino (SGSB) cluster emerged; at *K* = 9, the Klamath–Cascades (KC) cluster emerged; and at *K* = 10, the Central Coast South (CC-S) cluster separated from Central Coast Central (CC-C; **Fig. 3**). We observed no significant evidence for outlier loci using the Benjamini–Hochberg statistical correction in *tess3r* nor *BayeScan* for either *K* = 4 or *K* = 10.

**Figure 3.**
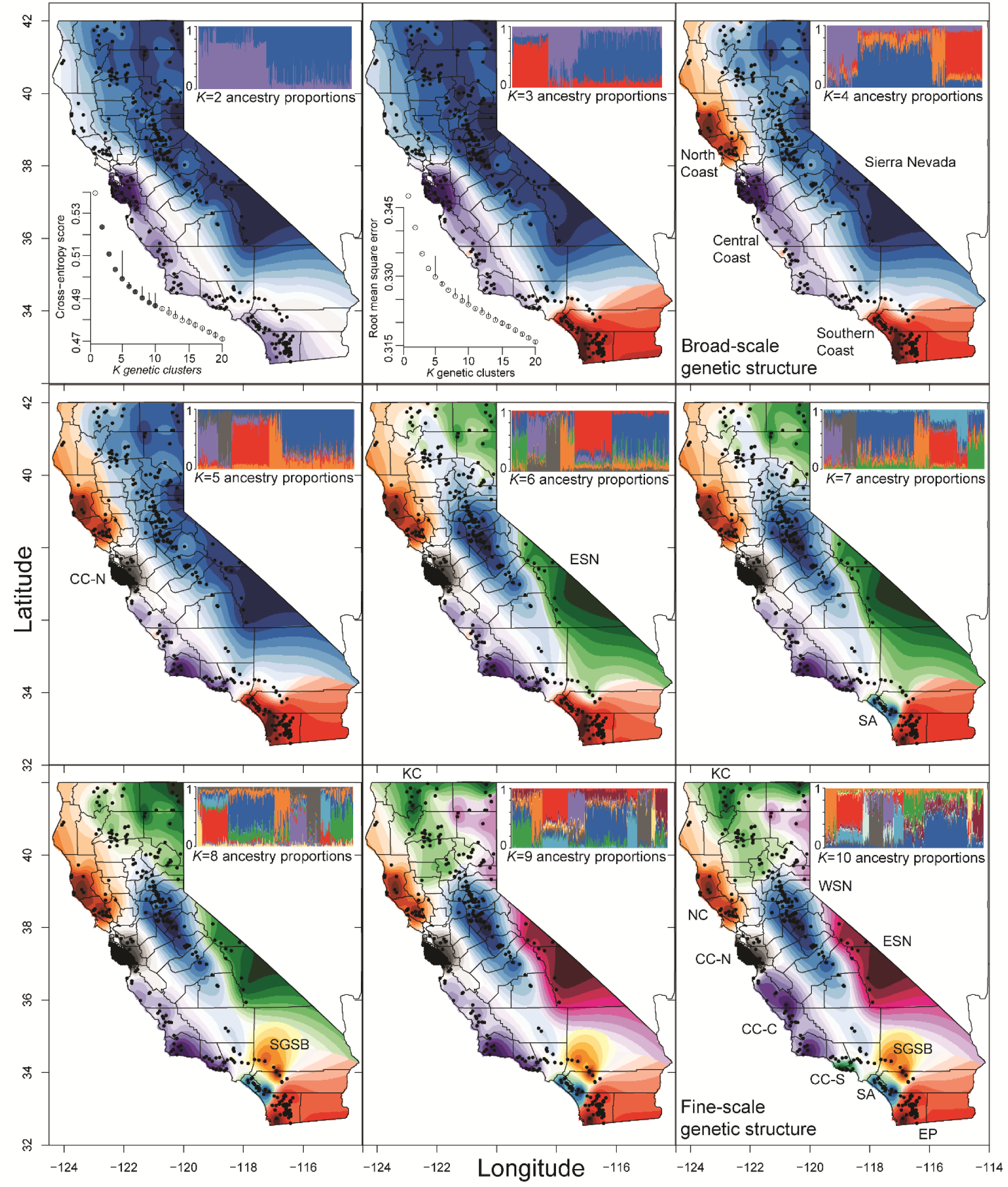
Interpolated ancestry proportions from *tess3r*, demonstrating the geographic distribution of biologically meaningful genetic clusters (*K*) ranging from 2–10. The “best” iterations of each *K*, based on cross-entropy score, is presented (shaded circles of inset plot in *K* = 2 panel). Root mean square error is also presented (inset plot in *K* = 3 panel). Both *tess3r* and the PCA (Fig. 2) support *K* = 4 and therefore the genetic clusters are labeled. At *K* = 10, fine-scale genetic clusters are labeled consistent with previous microsatellite data (Gustafson et al. 2019). For visualization, at each *K*, the genetic cluster that emerges is labeled. In alphabetical order, acronyms include Central Coast Central (CC-C), Central Coast North (CC-N), Central Coast South (CC-S), Eastern Peninsular Range (EP), Eastern Sierra Nevada (ESN), Klamath– Cascades (KC), North Coast (NC), Santa Ana (SA), San Gabriel–San Bernardino (SGSB), and Western Sierra Nevada (WSN).

### Genetic diversity, effective population size, genetic differentiation, and linkage decay

For *K* = 4, calculations of observed heterozygosity (*H*_O_), gene diversity (*H*_S_), polymorphic loci (*Poly*), allelic richness (*A*_r_), and the private alleles (*A*_p_) indicate that the Sierra Nevada cluster had higher genetic diversity than the Southern Coast, Central Coast, and North Coast (**Table 1**). Although significant, the North Coast was the only broad-scale genetic cluster that did not exhibit a strong Wahlund effect (i.e., significantly lower *H*_*O*_ compared to *H*_*S*_; SN: *t*=-50.6, *p*<0.001; SC: *t*=-48.2, *p*<0.001; CC: *t*=-58.5, *p*<0.001; NC: *t*=-10.6, *p*<0.001) or finer-scale substructure. Effective population sizes were not reported for broad-scale clusters because substructure introduced major biases (i.e., near-zero values) into *N*_e_ estimates.

**Table 1.** Heat map of genetic diversity statistics for *K* = 4 broad-scale and *K* = 10 fine-scale genetic clusters, including sample size (N), observed heterozygosity (*H*_*O*_); gene diversity (*H*_*s*_), proportion of polymorphic loci out of 16,285 (*Poly*), allelic richness corrected for sample size (*A*_*r*_), private alleles (*A*_*p*_), and effective population size (*N*_*e*_). Values for *N*_*e*_ are not presented for the *K* = 4 Sierra Nevada (SN), Southern Coast (SC), Central Coast (CC), or North Coast (NC) because of model assumption violations. There were no private alleles at *K* = 10, including Klamath–Cascades (KC), Western Sierra Nevada (WSN), Eastern Sierra Nevada (ESN), Eastern Peninsular (EP), San Gabriel–San Bernardino (SGSB), Santa Ana (SA), Central Coast Central (CC-C), Central Coast South (CC-S), Central Coast North (CC-N), and North Coast (NC).

| Genetic diversity |  | Genetic Cluster |  |  |  |  |  |  |  |  |  |
| --- | --- | --- | --- | --- | --- | --- | --- | --- | --- | --- | --- |
| $K = 4$ | | SN | | SC | | CC | | NC | | | |
| | $N$ | 193 | | 96 | | 79 | | 33 | | | |
| | $H_o$ | 0.31 | | 0.26 | | 0.24 | | 0.26 | | | |
| | $H_s$ | 0.34 | | 0.29 | | 0.28 | | 0.27 | | | |
| | $Poly$ | 0.93 | | 0.79 | | 0.77 | | 0.78 | | | |
| | $A_r$ | 1.79 | | 1.69 | | 1.67 | | 1.67 | | | |
| | $A_p$ | 37 | | 34 | | 17 | | 0 | | | |
| $K = 10$ | | KC | WSN | ESN | EP | SGSB | SA | CC-C | CC-S | CC-N | NC |
| | $N$ | 53 | 110 | 27 | 66 | 13 | 25 | 27 | 17 | 35 | 28 |
| | $H_o$ | 0.32 | 0.31 | 0.31 | 0.27 | 0.29 | 0.23 | 0.27 | 0.24 | 0.22 | 0.25 |
| | $H_s$ | 0.33 | 0.33 | 0.33 | 0.29 | 0.30 | 0.24 | 0.29 | 0.27 | 0.24 | 0.26 |
| | $Poly$ | 0.91 | 0.90 | 0.87 | 0.78 | 0.78 | 0.64 | 0.77 | 0.70 | 0.63 | 0.74 |
| | $A_r$ | 1.33 | 1.33 | 1.33 | 1.29 | 1.30 | 1.24 | 1.29 | 1.27 | 1.24 | 1.26 |
| | $N_e$ | 28.9 | 54.4 | 42.2 | 14.8 | 2.3 | 3.5 | 26.9 | 4.1 | 19.0 | 14.1 |

Broad-scale genetic clusters were moderately differentiated based on *F*_*ST*_ estimates which ranged from ∼0.1–0.2 (**Table 2**). The Sierra Nevada cluster was least differentiated from the others and the lowest *F*_*ST*_ estimates were between the Sierra Nevada and the North Coast clusters. In contrast, the Southern Coast cluster was the most differentiated from the others and the highest *F*_ST_ estimates were between the Southern Coast and the North Coast, followed by the Southern Coast and the Central Coast. At the broad scale, the linkage decay plot indicated that linkage disequilibrium (LD) was lowest in the Sierra Nevada and slightly increased in the Central Coast, Southern Coast, and North Coast clusters (**Fig. 4A**). When ignoring population status, California pumas (N = 401) had a LD *R*^*2*^ of ∼0.3 which decreased rapidly to less than 0.1 at a distance of 0.3 Mbp, then approached 0 at farther distances. Nearly the same result was observed in the Sierra Nevada. The Central Coast also had a major reduction in LD with distance but did not get under 0.1 until ∼3 Mbp in distance. In contrast, the Southern Coast and North Coast started with an LD *R*^*2*^ of ∼0.4 which remained above 0.1 even at distance of 8 million bp (**Fig. 4A**).

**Table 2.** Heat map of mean pairwise genetic distance values for the broad-scale *K* = 4 and fine-scale *K* = 10 genetic clusters. Weir and Cockerham *F*_*ST*_ is presented below the diagonal and Nei’s *F*_*ST*_ is presented above the diagonal (WC\Nei). All pairwise estimates were significant (*p* < 0.001) based on a bootstrapping analysis using *hierfstat::boot*.*ppfst*. Sierra Nevada (SN), Southern Coast (SC), Central Coast (CC), North Coast (NC), Klamath–Cascades (KC), Western Sierra Nevada (WSN), Eastern Sierra Nevada (ESN), Eastern Peninsular (EP), San Gabriel–San Bernardino (SGSB), Santa Ana (SA), Central Coast Central (CC-C), Central Coast South (CC-S), Central Coast North (CC-N).

| WC\Nei $F_{ST}$ | | Genetic Cluster | | | | | | | | | |
| --- | --- | --- | --- | --- | --- | --- | --- | --- | --- | --- | --- |
| $K=4$ | | SN | | SC | | CC | | NC | | | |
|  | SN | - |  | 0.133 |  | 0.124 |  | 0.100 |  |  |  |
|  | SC | 0.129 |  | - |  | 0.173 |  | 0.198 |  |  |  |
|  | CC | 0.120 |  | 0.173 |  | - |  | 0.156 |  |  |  |
|  | NC | 0.094 |  | 0.196 |  | 0.156 |  | - |  |  |  |
| $K=10$ | | KC | WSN | ESN | EP | SGSB | SA | CC-C | CC-S | CC-N | NC |
|  | KC | - | 0.022 | 0.041 | 0.141 | 0.109 | 0.215 | 0.117 | 0.146 | 0.183 | 0.093 |
|  | WSN | 0.022 | - | 0.045 | 0.149 | 0.111 | 0.222 | 0.121 | 0.147 | 0.188 | 0.126 |
|  | ESN | 0.041 | 0.045 | - | 0.163 | 0.116 | 0.226 | 0.168 | 0.189 | 0.233 | 0.183 |
|  | EP | 0.141 | 0.146 | 0.166 | - | 0.130 | 0.100 | 0.164 | 0.196 | 0.231 | 0.214 |
|  | SGSB | 0.105 | 0.106 | 0.113 | 0.132 | - | 0.212 | 0.140 | 0.163 | 0.210 | 0.205 |
|  | SA | 0.202 | 0.203 | 0.221 | 0.095 | 0.217 | - | 0.254 | 0.2869 | 0.3198 | 0.3013 |
|  | CC-C | 0.114 | 0.116 | 0.168 | 0.163 | 0.141 | 0.251 | - | 0.060 | 0.098 | 0.164 |
|  | CC-S | 0.137 | 0.136 | 0.183 | 0.192 | 0.164 | 0.289 | 0.059 | - | 0.148 | 0.202 |
|  | CC-N | 0.178 | 0.176 | 0.237 | 0.227 | 0.221 | 0.320 | 0.100 | 0.152 | - | 0.229 |
|  | NC | 0.090 | 0.118 | 0.183 | 0.211 | 0.210 | 0.300 | 0.164 | 0.203 | 0.230 | - |

**Figure 4.**
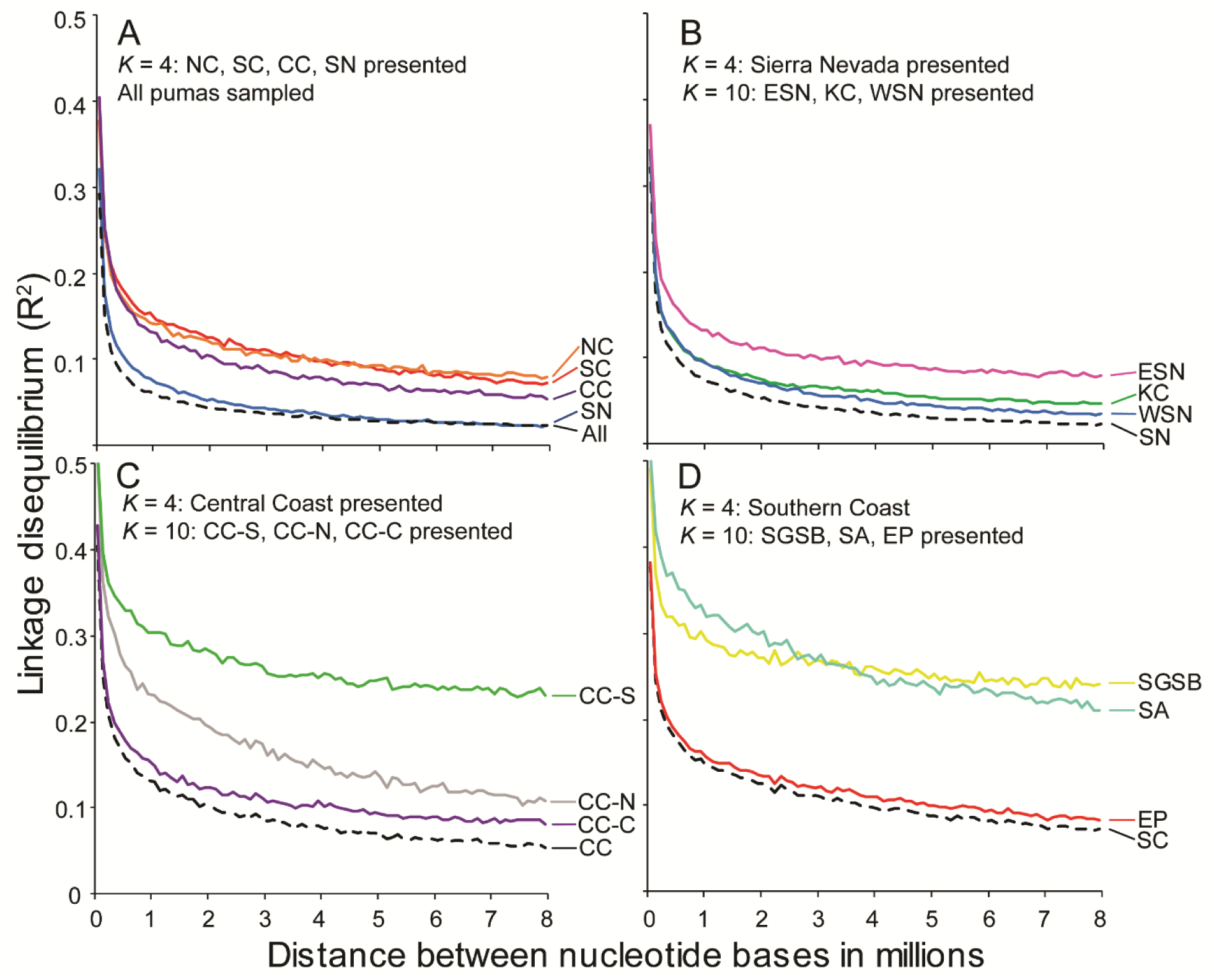
Correlation of SNPs with genomic distance, ranging from hundreds to 8 million nucleotides in distance. Based on pairwise estimates from 16,285 SNPs, linkage decay is presented for all 401 pumas sampled in California (All), from the *K* = 4 broad-scale genetic clusters (**A**: North Coast, NC; Southern Coast, SC; Central Coast, CC; Sierra Nevada, SN), and from the *K* = 10 fine-scale genetic clusters (**B–D**). Fine-scale clusters are presented within their corresponding broad-scale group. The NC is only presented in the first panel because it did not exhibit substructure. (**B**) Eastern Sierra Nevada (ESN), Klamath–Cascades (KC) and Western Sierra Nevada (WSN) are nested within SN. (**C**) Central Coast South (CC-S), Central Coast North (CC-N), and Central Coast Central (CC-C) are nested within CC. (**D**) San Gabriel–San Bernardino (SGSB), Santa Ana (SA), and Eastern Peninsular Range (EP) are nested within SC. In each figure, the dashed line represents the broadest-scale designation within the group.

Fine-scale genetic clusters within the Sierra Nevada — including KC, WSN, and ESN — had the highest genetic diversity estimates, as well as the highest estimates of *N*_*e*_. Only the WSN had an *N*_*e*_ above 50, a threshold commonly considered to be sustainable over the long-term (**Table 1**; Franklin, 1980). Pairwise *F*_ST_ estimates among fine-scale genetic clusters within the Sierra Nevada suggested weak substructure, with little genetic differentiation (i.e., pairwise *F*_*ST*_ < 0.05), indicating substantial gene flow throughout this region (**Table 2**). Within the Sierra Nevada, the ESN showed slightly higher LD than KC or WSN, and all three retained a high proportion of polymorphic loci.

The fine-scale genetic clusters within the Southern Coast — including EP, SGSB, and the SA — exhibited lower genetic diversity estimates when compared to the Sierra Nevada, as well as large differences when compared to each other (**Table 1**). Estimates were generally lowest in SA, whereas EP and SGSB had similar overall estimates. However, both SA and SGSB had extremely low estimates of *N*_*e*_. Unlike the Sierra Nevada, fine-scale genetic clusters within the Southern Coast had moderate to strong genetic differentiation from one another (pairwise *F*_*ST*_ values ∼0.1–0.2; **Table 2**). Except for the moderate differentiation with EP (i.e., pairwise *F*_*ST*_ of ∼0.1), SA was the most differentiated among the 10 fine-scale genetic clusters (pairwise *F*_*ST*_ values range: ∼0.2–0.3). The SGSB cluster had relatively lower pairwise *F*_*ST*_ estimates with the Sierra Nevada and EP clusters, moderate *F*_*ST*_ estimates with CC-C and CC-S, and was more strongly differentiated from the CC-N and NC. The EP cluster showed similar patterns of differentiation but was least differentiated from the geographically adjacent SA and SGSB clusters. Although EP exhibited LD estimates similar to the Southern Coast as a whole, SGSB and SA started with a high LD *R*^*2*^ of ∼0.5 which decreased to just above 0.3 at a distance of 0.3 Mbp, then remained high (above 0.25) at farther distances (**Fig. 4**).

The fine-scale genetic clusters within the Central Coast exhibited the most variation in estimates of genetic diversity (**Table 1**). The CC-C cluster had the highest diversity within the region, including the largest estimate of *N*_*e*_. The CC-S cluster had intermediate levels of diversity but exhibited the lowest *N*_*e*_ in the region. The CC-N cluster had as low, or lower, genetic diversity estimates than most of the 10 fine-scale genetic clusters examined overall, but had one of the higher *N*_*e*_ estimates outside of the Sierra Nevada. Differentiation within the Central Coast was moderate overall (pairwise *F*_ST_ ∼0.06–0.15) and appeared to correlate with distance (i.e., CC-N more differentiated from CC-S than CC-C; **Table 2**). Within the Central Coast, CC-C had the lowest LD *R*^*2*^ values (**Fig. 4**). The CC-N cluster had higher LD values, especially at lower distances between SNPs, whereas CC-S had among the highest LD R^2^ values, comparable to those of SGSB and SA in the Southern Coast.

Finally, the NC had genetic diversity estimates that were lower than the Sierra Nevada and comparable to the Southern Coast and Central Coast, with an *N*_*e*_ estimate of 14.1 (**Table 1**). Overall, the NC showed strong differentiation from the other fine-scale genetic clusters with the exception of KC and WSN for which differentiation was moderate (**Table 2**). The linkage decay plot indicates the NC had similar LD *R*^*2*^ values to that of ESN and EP (**Fig. 4**).

## DISCUSSION

Our analyses of genetic diversity and linkage disequilibrium on 16,285 SNPs from 401 pumas throughout California demonstrated that the complex geography and land use patterns in California result in equally complex patterns of gene flow and population structure. The high-density SNP data provided resolution to detect both four broad-scale genetic clusters with high genetic diversity, as well as substructure that we designate as 10 fine-scale genetic populations with highly variable genetic diversity. Our data further supports the notion that puma populations in California form a “horseshoe” network around the Central Valley with San Francisco Bay acting as a barrier to gene flow along the coast (Gustafson et al., 2019). For the Sierra Nevada cluster, the nested fine-scale populations had consistently high genetic variation. However, within the coastal groups, genetic variation within certain fine-scale genetic populations was concerningly low, while others appeared to have retained sufficient variation as to be capable of serving as sources of genetic rescue under various management scenarios (i.e., assisted migration). In fact, our linkage decay analysis indicated that populations with low genetic diversity and high linkage disequilibrium do not necessarily share the same fixed loci, consistent with what was suggested by Saremi et al. (2019). Therefore, maintaining and enhancing connectivity within and among broad-scale groups could increase genetic diversity to entire regions and could decrease the apparent effects of genetic drift and inbreeding to some at-risk coastal populations (Ernest et al., 2003; 2014; Gustafson et al., 2017; Riley et al., 2014).

The evidence for four broad-scale genetic groups is different than previous studies using microsatellites (Ernest et al. 2003; Gustafson et al., 2019) indicating the importance of using genomic methods in the study of broader-scale wildlife conservation genetics. Our data further supports the claim that the Sierra Nevada region is a major refugium of puma genetic diversity in California (Gustafson et al., 2019). Therefore, it is important to protect the Sierra Nevada group from habitat degradation and foster conservation actions that can enhance gene flow with the North Coast, Central Coast, and Southern Coast clusters as well as the Great Basin to the east (Gustafson et al., 2019). The broad-scale Southern Coast group is least connected to the other genetic clusters in the state but had higher genetic diversity and more private alleles than the Central Coast or North Coast. This indicates that the Southern Coast group retains unique genomic variation that must be conserved in order to maximize genetic diversity among pumas in California. Although the four major genetic clusters are highly consistent among our structure analysis and PCA, there was also statistical support for substructure (i.e., *tess3r* results and moderate to high pairwise F_ST_ values within and among the broad-scale groups) indicating as many as 10 fine-scale genetic populations.

Generally, the 10 genetic populations identified with SNPs correspond strongly to those identified in previous studies using microsatellite markers and different samples (Ernest et al., 2003, Gustafson et al., 2019). However, the northern-most Klamath–Cascade population was not observed previously with microsatellites (Gustafson et al., 2019). This is likely because there were very few pumas available for analysis in the Klamath or Cascade Mountains during the 2019 microsatellite study. It is also possible that 42 microsatellites may not have been sufficient to detect the low genetic differentiation (*F*_*ST*_ = 0.022) observed between the Klamath–Cascade and Western Sierra Nevada populations. The 10 populations varied considerably in genetic diversity estimates (*H*_*O*_ range 0.22–0.32; *H*_*S*_ range 0.24–0.33; *Poly* range: 0.63–0.91; *A*_*r*_ range: 1.24–1.33), effective population sizes (*N*_*e*_ range 2.3–54.4), and genetic differentiation (*F*_*ST*_ range: 0.22–0.32), as discussed below.

A major difference between this and previous studies is the observation that pumas in the Central Coast North population have genetic diversity estimates as low as those in the Santa Ana and Central Coast South populations, which are highly isolated by urbanization and transportation infrastructure and exhibit evidence of inbreeding depression (Benson et al., 2020; Ernest et al., 2014; Gustafson et al. 2017; Riley et al., 2014; Vickers et al., 2015). Our results are consistent with those of Saremi et al. (2019), which indicated that inbreeding metrics between pumas from the Santa Monica Mountains (in Central Coast South) and pumas from the Santa Cruz Mountains (in Central Coast North) were similar. Interestingly, *N*_e_ for the Central Coast North was much higher than both the Santa Ana and Central Coast South populations. These observations are consistent with a large breeding population experiencing genetic drift due to dispersal barriers to the north (i.e., San Francisco Bay) and gene flow only occurring with the Central Coast Central population to the south. This pattern could also be driven by carrying capacity processes associated with habitat limitations (Dellinger, Gustafson, Gammons, Ernest, & Torres, 2020b). If dispersal is limited by continued development southeast of the Central Coast North population, rapid genetic drift and inbreeding may ensue (Mills & Allendorf, 1996; Wang, 2004) and local extinctions may occur as predicted in the Central Coast South and Santa Ana populations (Benson et al., 2016; 2019). Thus, puma population viability will be facilitated when land management agencies and land developers in the region work proactively to preserve or enhance wildlife corridors.

Notably, the San Gabriel–San Bernardino population had the lowest *N*_e_, but had intermediate levels of genetic diversity. Occasional migrants could alter *N*_*e*_ estimates and temporarily inflate estimates of heterozygosity (Gustafson et al., 2017). We suggest this could also be the result of metapopulation dynamics—i.e., a small local population with frequent turnover located at the intersection of dispersal corridors for the Sierra Nevada, Central Coast, and Southern Coast groups. Although the genetics of this population are complex and somewhat uncertain, this region is of critical importance for maintaining statewide puma gene flow.Enhancing connectivity through the Transverse Ranges (including the Tehachapi Mountains, Sierra Pelona, San Gabriel Mountains, and San Bernardino Mountains; **Fig. 1B**) is a critical conservation priority in order to maintain gene flow between the Southern Coast populations and the Sierra Nevada or Central Coast groups.

The three populations with the lowest *N*_e_, including the San Gabriel–San Bernardino, Santa Ana, and Central Coast South populations, have the smallest available amount of habitat (Dellinger et al., 2020b), and had the highest linkage disequilibrium throughout their genomes. As we observed, there was great variation among populations in the decay curves, with the Central Coast North population having the next highest linkage disequilibrium after these three populations. Given the genetic diversity, *N*_*e*_, and linkage data, the San Gabriel–San Bernardino and Central Coast North populations may be approaching levels of genetic drift and inbreeding similar to the well-monitored and genetically depauperate Santa Ana and Central Coast South populations (Ernest et al., 2014; Gustafson et al., 2017; Riley et al. 2014).

Populations with intermediate genetic diversity include the North Coast, Central Coast Central, and Eastern Peninsular Range. Measures of genetic diversity were lower than expected for the North Coast population given there are no obvious anthropogenic barriers to gene flow with the Klamath–Cascade, Western Sierra Nevada, or pumas from Oregon (Gustafson et al., 2019). However, the majority of our samples from this genetic cluster came from just north of the San Francisco Bay, an area of substantial human density and an area surrounded by ocean or the Central Valley. Thus our results may not be truly representative of this region as a whole and may represent the most isolated pumas in the “peninsula” of habitat. Future studies would benefit from increased sampling throughout this genetic cluster, north to (and including) Oregon. Nonetheless, pumas and other animals would benefit if decisions for future development between the North Coast and Sierra Nevada consider the future connectivity of private timber land holdings along the coast with the inland National Forests.

The Central Coast Central population has ample habitat for maintaining a breeding population (Dellinger et al., 2020b). Given the apparent absence of gene flow across the Central Valley, this population may be the only consistent source of migrants for the Central Coast North and Central Coast South, which have concerningly low levels of genetic diversity and evidence of inbreeding. Thus, we consider the Central Coast Central population to be essential for the long-term viability of both adjacent populations, and urge that habitat in this region not be fragmented further.

Despite having less than half of the overall habitat of the Central Coast Central population (Dellinger et al., 2020b), the Eastern Peninsular Range population has roughly similar genetic diversity estimates, but a much lower *N*_e_. Dispersal in and out of the Eastern Peninsular Range is extremely limited and the degree to which pumas disperse across the border between USA and Mexico remains unknown (Gustafson et al., 2019). Given that the Eastern Peninsular Range is the only population known to exchange individuals with the Santa Ana population, management actions which enhance gene flow between these areas remain critical to the recovery of pumas in the Santa Ana Mountains.

Our linkage decay analysis indicated that in the Central Coast South, San Gabriel–San Bernardino, Santa Ana, and perhaps the Central Coast North populations, pumas have long runs of homozygosity that are identical-by-descent. This finding is consistent with the genome resequencing results of Saremi et al. (2019) in the Santa Cruz (i.e., Central Coast North) and Santa Monica Mountains (i.e., Central Coast South), which suggested close and recent inbreeding led to runs of homozygosity. Although Saremi et al. (2019) only sequenced individuals from California populations known to have low genetic diversity, our linkage decay results from populations throughout the state indicate that the genome-level problems of inbreeding and homozygosity are not universal throughout California. Instead, the Klamath– Cascades, Western Sierra Nevada, Eastern Sierra Nevada, Central Coast Central, and the Eastern Peninsular Range populations all have low linkage disequilibrium throughout the genome. Additionally, when the inbred populations are analyzed with their broad-scale group, linkage decay curves demonstrated the potential for remaining diversity to reduce linkage to negligible levels through gene flow and meiotic recombination. We observed up to 30–37% of the SNPs as fixed in the Central Coast–South, Santa Ana, and Central Coast North populations. Our linkage decay curves and the genome resequencing results of Saremi et al. (2019) in two of these areas demonstrates that the fixations are different among populations. Thus, genetic restoration is possible even among genetically depauperate populations. When considering that genetic diversity is much higher in several California puma populations than those heavily studied along urban coasts, there is high potential for the long-term persistence of pumas in throughout the majority of California.

We tested for outlier loci using multiple methods (Narum & Hess, 2011) but found no evidence of local adaptation when *K* = 4 or *K* = 10. Detection of outlier loci with RADseq is limited by the reduced representation of the genome, yet it has often been shown to be an effective approach (Catchen et al., 2017). Pumas are long-distance dispersers (Hawley et al., 2016; Sweanor, Logan, & Hornocker, 2000) and inhabit all major mountain ranges in California (Dellinger et al., 2020b), suggesting local adaptation may be unlikely. Our results provide preliminary evidence that outbreeding depression resulting from potential active genetic management may be of minimal concern (Frankham et al., 2011). Recent modelling (Kyriazis, Wayne, & Lohmueller, 2021) does suggest, however, that attempts to maximize genetic diversity in a population can introduce hidden deleterious recessive mutations, enhancing extinction risk. The modelling of Kyriazis et al. (2021) has faced criticisms (Garcia-Dorado & Caballero, 2021), however, and Ralls, Sunnucks, Lacy, and Frankham (2020) argue that the benefits of increasing genetic diversity outweigh the risks. Thus, managers could consider assisted migration (e.g., wildlife overpasses/underpasses, translocation of individuals between populations, etc.) to improve viability of some coastal populations, as was empirically demonstrated to have shifted the trajectory of Florida panther population from extinction (Ralls et al., 2018). However, we suggest whole genome resequencing methods better suited for detecting selection (Fuentes-Pardo & Ruzzante, 2017) be implemented before such efforts, especially over long distances. Managers would also need to consider other risks as well, such as the movement of pathogens or the ethical implications of moving large carnivores (Bevins et al., 2012). Wildlife managers will have to weigh these concerns against their obligation to minimize the risks of extirpation such as those predicted for the Santa Ana and Central Coast South populations (Benson et al., 2019), and shown here to be a concern in the Central Coast North population as well. If assisted migration is contemplated, then these factors, as well as possible local adaptation, should be weighed carefully. It is our opinion that current efforts to construct or improve wildlife overpasses that can facilitate natural migration into coastal populations should be considered the primary management strategy for conserving viable puma populations in that region.

## CONCLUSION

Our hierarchical population genomic analyses provide decision makers a contemporary and thorough evaluation of the genetic diversity, effective population sizes, and connectivity of puma populations throughout California. These state- and genome-wide results are critically important for conservation and management practices in California, especially considering the increasing demand for development and the current political climate surrounding the status of the puma (Yap & Rose, 2019). In brief, puma populations are widespread throughout the mountains of California. Populations range from major genetic sources to populations with issues of low genetic diversity and inbreeding. The inbred populations do not share the same runs of homozygosity and therefore genetic diversity could be restored through enhanced gene flow. Current challenges to puma populations are highly regional and should be addressed by focusing on how natural geography and human development impacts puma habitat and movements locally. Attention is understandably given to those populations that are highly imperiled, but it is important to note that California has several thriving populations throughout the state which represent an important resource for any genetic management strategy. Protecting tracts of contiguous habitat to preserve large populations will provide greater protection for the species as a whole. Specifically, further fragmentation of habitat in the Sierra Nevada group could be catastrophic to population viability of pumas in the state because it serves as a genetic refugium. Protecting, enhancing, and creating movement corridors to allow statewide “stepping-stone” connectivity at broad and fine scales will allow for the migrants needed to counteract the local extirpations faced by some coastal populations.

## ACKNOWLEDGMENTS

We thank the multiple agencies and people who provided samples and expertise including California Department of Fish and Wildlife (D. Clifford, R. Botta, J. Colby, S. Torres), California State Parks, The Nature Conservancy, University of California Davis Wildlife Health Center (T. Drazenovich), University of California Los Angeles, The National Park Service, and the US Geological Survey. We thank University of Wyoming personnel, S. Love Stowell, A. Gustafson, L. Johnson, B. Godwin, and C. A. Buerkle. Computational resources were provided by Advanced Research Computing Center (2018) Teton Computing Environment, Intel x86_64 cluster. University of Wyoming, Laramie, WY https://doi.org/10.15786/M2FY47. This research was also supported by the Arkansas High Performance Computing Center which is funded through multiple National Science Foundation grants and the Arkansas Economic Development Commission. Funding for this work was provided by the California Department of Fish and Wildlife (H.B.E.), Excellence Chair funds (H.B.E), National Science Foundation Ecology of Infectious Disease program grant (DEB 1413925).

## AUTHOR CONTRIBUTIONS

KDG, MRB, JAD, and HBE designed the study. KDG, RBG, JLR, and MEFL performed laboratory research. MRB, TWB, SPDR, JS, and JAD performed field research. KDG, RBG, MRB, MEFL, and HBE analyzed the data. KDG, RBG, MRB, TWV, SPDR, JS, JLR, JAD, MEFL, and HBE wrote the paper.

